# Overcoming the dichotomy: new insights into the genomic diversity of open and isolated European populations

**DOI:** 10.1101/067850

**Authors:** Paolo Anagnostou, Valentina Dominici, Cinzia Battaggia, Luca Pagani, Miguel Vilar, Spencer Wells, Davide Pettener, Stefania Sarno, Alessio Boattini, Paolo Francalacci, Vincenza Colonna, Giuseppe Vona, Carla Calò, Giovanni Destro Bisol, Sergio Tofanelli

**Author notes:** These two authors contributed equally (Paolo Anagnostou, Valentina Dominici, Giovanni Destro Bisol, Sergio Tofanelli).

## Abstract

Human populations are often dichotomized into “isolated” and “open” using cultural and/or geographical barriers to gene flow as differential criteria. Although widespread, the use of these alternative categories could obscure further heterogeneity due to inter-population differences in effective size, growth rate, and timing or amount of gene flow. We compared intra and interpopulation variation measures combining novel and literature data relative to 87,818 autosomal SNPs in 14 open populations and 10 geographic and/or linguistic European isolates. Patterns of intra-population diversity were found to vary significantly more among isolates, probably due to differential levels of drift and inbreeding. The relatively large effective size estimated for some population isolates challenges the generalized view that they originate from small founding groups. Principal component scores based on measures of intra-population variation of isolated and open populations turned out to be distributed along a sort of continuum, with an area of intersection between the two groups. Patterns of inter-population diversity were even closer, as we were able to detect some differences between population groups only for a few multidimensional scaling dimensions. Therefore, different lines of evidence suggest that dichotomizing human populations into open and isolated fails to capture the actual relations among their genomic features.

## INTRODUCTION

Human groups that have been subject to geographical and/or socio-cultural barriers (e.g. linguistic, social or religious) to inward gene flow during their evolutionary history are commonly referred to as isolated or closed populations (hereafter “isolates/isolated”). In current genetic literature, they are often opposed to open or outbred populations - exempt from known limitations to admixture - since a higher level of inbreeding and drift and a lower efficiency of recombination in redistributing variation across individuals are to be expected under isolation^1,2,3^. Although common practice, the use of the terms *open* and *isolated* to indicate two discrete dichotomous categories could obscure the existence of further heterogeneity. In fact, when applying such a distinction to genetics based on environmental and socio-cultural factors, we implicitly assume that it is not confounded by inter-population differences in effective size, growth rate and timing or extent of gene flow reduction. Unfortunately, current knowledge on human isolates cannot help us understand whether this is consistent with the patterning of genetic diversity.

A number of studies has investigated the effects of isolation in human populations by exploiting the high sensitivity to drift of unilinear markers of mtDNA and the non-recombining portion of the Y chromosome^4,5,6,7,8,9^. However, the lack of recombination limits the power of these genetic systems in the detection of signatures of genetic isolation in different historical and demographic conditions.

With the introduction of SNP microarrays, which enable the simultaneous analysis of hundreds of thousands of loci distributed across the genome, it is now possible to investigate the genetic structure of human populations on a fine scale^10^. Using autosomal variation, we can detect signatures of isolation which are not revealed by unilinear markers, such as the increase in the number and size of stretches of consecutive homozygous genotypes, shared chromosomal segments identical by descent (IBD) and Linkage Disequilibrium (LD)^11,12,13,14,2,3^. Investigations published so far have provided accurate genetic characterizations of a number of human genetic isolates, with a prevalent focus on one or few populations and their potential use in gene-disease association studies^15,16,17,18,19^. Relations between genomic differences and demographic or historical factors and their implications for the gene mapping of Mendelian or complex traits were also studied^1,2,3^, while LD patterns were compared in isolates distributed worldwide^14^. More recently, other studies have simultaneously investigated multiple isolates, mostly focusing on populations with shared historical and demographic features^17,18,20^. However, to the best of our knowledge, no study has systematically explored the structure of genomic diversity in isolated populations comparing them with a comprehensive set of open populations. The European continent provides optimal conditions for these investigations. There is, in fact, broad convergence regarding the notion that European genomic diversity has been shaped primarily by geography, with the isolation-by-distance model being well supported even at long latitudinal distances^21,22,23^. Therefore, the comparative study of open and isolated populations may be performed in wider transects with less confounding factors than in other continental areas.

Here we present a study of 24 European populations, nine of which were newly genotyped using the GenoChip 2.0 array^24^. Our dataset includes ten populations, which have been subject to linguistic and/or geographic barriers to gene flow during their recent history. We compare their intra- and inter-population measures of variation in order to evaluate the apportionment of genomic diversity in the populations under study, and to better understand to what extent the discrete open and isolated dichotomous categories correspond to the way in which the genome is structured.

## RESULTS

### Genomic variation in open and isolated European populations

In order to better understand how genomic diversity is structured in open and isolated populations, we first analysed the distribution of seven intra-population measures of single nucleotide (homozygosity, interlocus variance and intra-pop pairwise IBS) and haplotype variation (average number and total length of RoHs, average IBD block sharing and length of LD blocks). As expected, their median values were significantly higher in isolates (alpha=0.01, Mann-Whitney U test), with the length of linkage blocks being the only exception (Fig. 2). However, an overlap between the two groups was observed for six out of seven measures. The most evident one was shown by the average length of LD blocks, in which seven isolates fall within the range of open populations. A clear-cut distinction between the two groups was provided by intra-population IBS pairwise identities only. Similarly, variance between populations was higher among isolates for six measures, and the difference turned out to be statistically significant for four of them (Levene Test, Fig. 2). The largest one was observed for the average intra-population IBD block sharing, whose standard deviation was found to be 28 times higher in isolated populations. We also investigated the distribution of RoHs in more detail, because their length and number have been shown to vary between open and isolated populations^32,40,41^. Although isolated populations had a significantly higher number of RoHs, an overlap between the two groups was observed for all size classes (Supplementary Figure S2).

**Figure 2.**
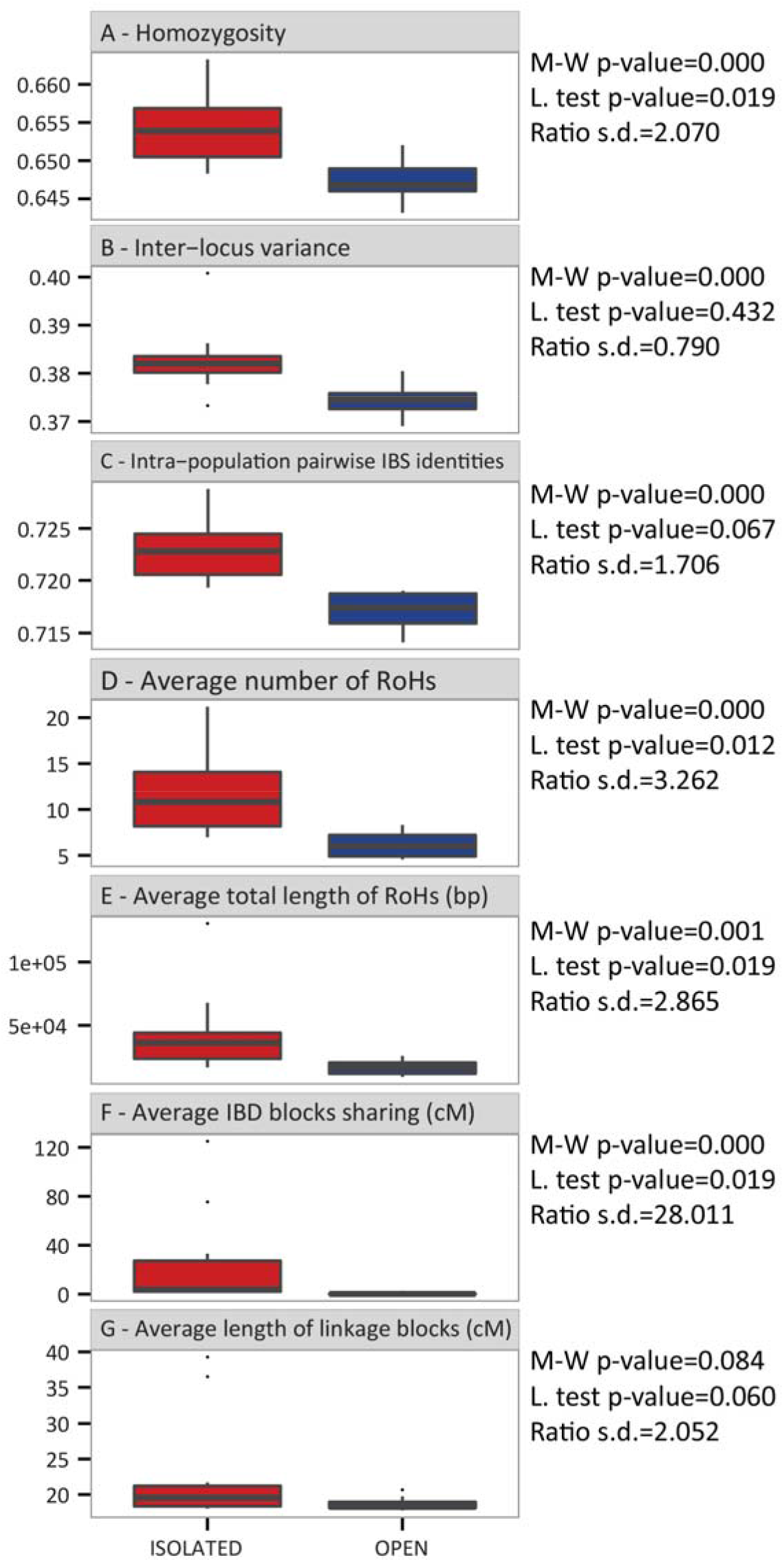
Boxplots of (A) Average homozygosity over loci (B) Inter-locus variance (C) Average number of RoHs (D) Average total length of RoHs (E) Average intra-population pairwise IBS (F) Average population IBD blocks sharing (G) Average length of linkage blocks. M-W stands for Mann-Whitney U test, L. for Levene test and s.d for standard deviation (these last two tests were performed excluding the outliers).

Moving on from groups to single populations, we observed the strongest signals of isolation in Sauris and Sappada (Supplementary Figures S3 and S4). Due to its small sample size (N=10; see Supplementary Table S1), the evidence for the former population was tested with resampling procedures, obtaining consistent results. These two populations also have the highest proportion of long RoHs (classes 5 and 6), whereas Basques and Benetutti prevail for the small and medium ones (classes 1 to 3) (Supplementary Figure S5). By contrast, the weakest signal of isolation is provided by Cimbrians, who show the lowest values for all measures. Inter-individual variation reached the highest values in Sauris, Timau and Sappada, whose values were found to be significantly higher in at least 70% of pairwise comparisons with open populations (see Supplementary Tables S2-S6). A less intense but noticeable signal was observed for North Sardinia, Benetutti and the Basques, which are the only remaining populations with a proportion of significant pairwise comparisons above 50%.

We combined all the measures of intra-population variation using a PCA to obtain a synthetic view of signatures of isolation (Fig. 3A). All variables heavily load on the first component, which describes 69.7% of the total variance, with the highest contributions by RoHs (total number and length; proportion of medium and large RoHs), intra-population IBS and Homozygosity (see Fig. 3B). Overall, the first component separates isolates (on the left) from open populations (on the right), with the German-speaking island of Sauris and the Bulgarians at the two extremes. The second principal component, which describes 17.2% of the total variance, does not set open and isolated populations apart, although the former are more tightly clustered. Sauris (at the upper side), Basques, and Benetutti (at the lower side) are found at the poles of the distribution. Among the factors that contribute most to the positive scores are the average values for the number of very long RoHs (class 6), length of LD block and population IBD block sharing, whereas the average number of small and intermediate RoH (class 1 to 3) load on negative scores. When using more relaxed settings for the RoH identification (minimum number of SNP=12), no substantial difference was observed for the population distribution of number and total size of RoHs, proportion of the RoH classes and the resulting PCA plot (Supplementary Table S7).

**Figure 3.**
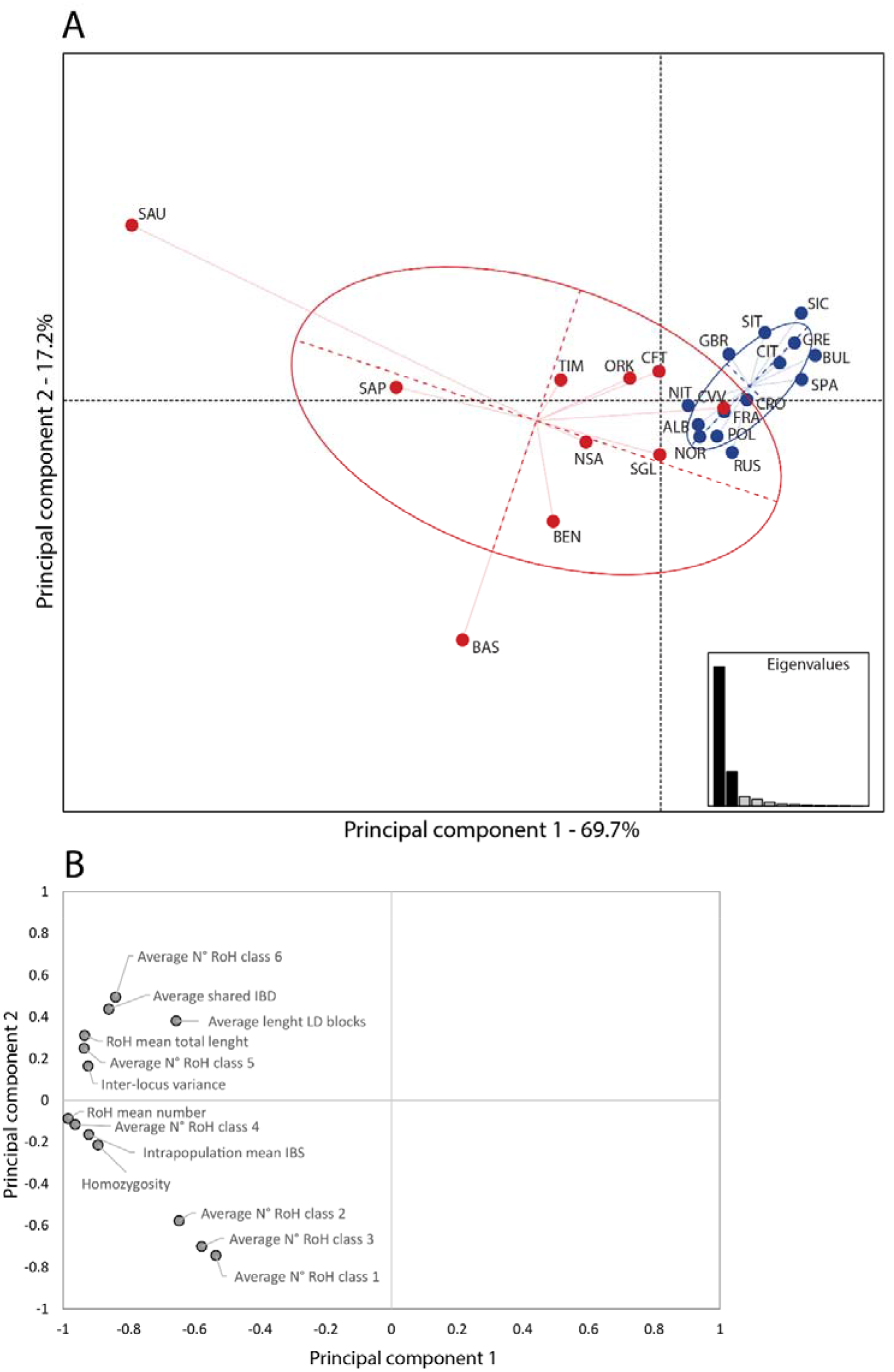
Principal component plots based on intra-population measures. (A) Scatter plot of the first two principal components. (B) Plot of the factor scores for the first and second principal components. Labels as in Table 1.

### Effective population size in open and isolated European populations

Among isolated populations, the Ne values range between 209 (Sauris, 208–210; 95% confidence interval) and 3739 (Basque, 3607–3880; 95% confidence interval; Fig. 4, Supplementary Table S8). Regarding open populations, the values range between 2386 (Albania, 2342–2452; 95% confidence interval) and 8267 (Poland, 7850–8732; 95% confidence interval). Noticeably, seven open (Albania, Croatia, Greece, Aosta, Sicily, South Italy and Norway) and five isolated populations (Basques, Benetutti, Carloforte, North Sardinia, and Cimbrians) fall into the range of the alternative group. When repeating the estimates in the four populations for which SNP data at a higher resolution were available (HGDP panel; 647,789 SNP) using an IBD sharing based method^42^, we obtained values that were different in absolute terms but highly correlated with those produced by GenoChip 2.0 (see Supplementary Table S9).

**Figure 4.**
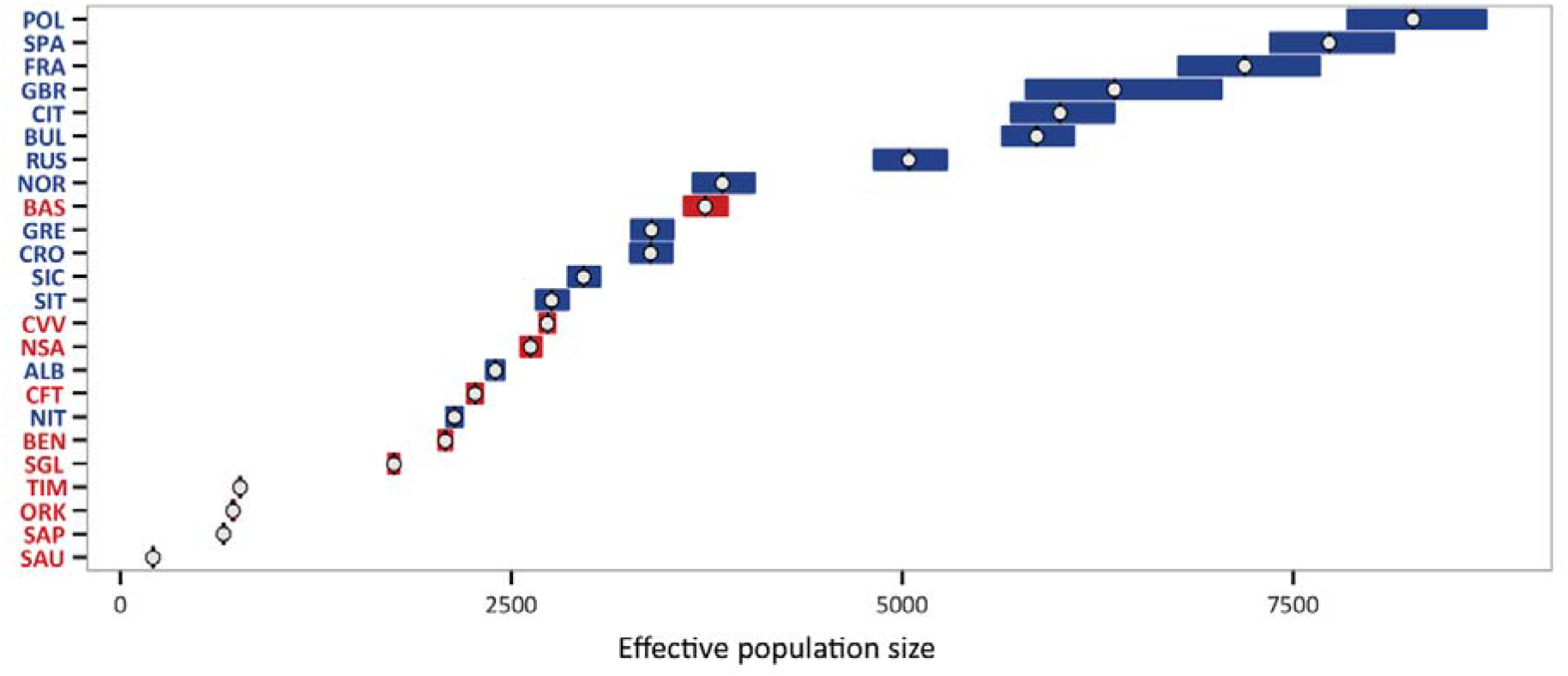
Effective population size estimates based on 68,205 SNPs (16 chromosomes). White circles and bars represent point estimates and 95% confidence interval, respectively. Abbreviations as in Table 1.

Fig. 5 displays a range of possible combinations of N_e_ and time since isolation (coloured areas) able to produce the inbreeding coefficients observed in four isolated populations (see Materials and Methods), with an indication of time since their foundation as suggested from historical and genetic sources (Supplementary Text S1). At any given N_e_, the values for time since isolation relative to Basques and North Sardinians were found to be lower than in Sappada and Sauris. The ratio ranged from approximately two to three times lower compared to Sappada and four to eight times higher compared to Sauris (Supplementary Table 10).

**Figure 5.**
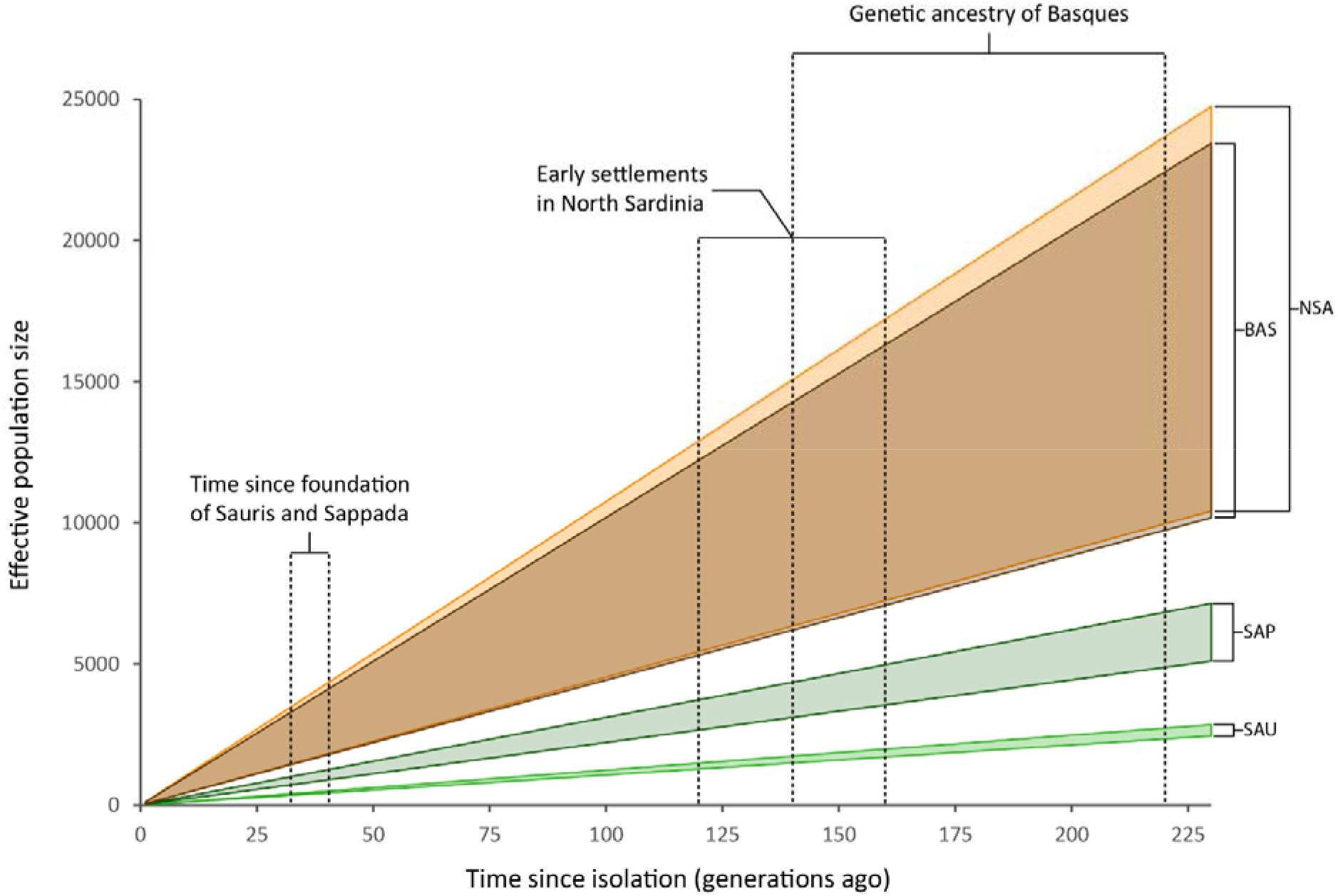
Numbers of generations since isolation (X axis) and corresponding N_e_ values (Y axis) under a model of constant population size in populations retaining clear signatures of isolation (Basques, North Sardinia, Sappada and Sauris; see also PC1 in Fig. 3). For each population at any given time since isolation, upper and lower boundaries of Ne were obtained assuming the initial inbreeding coefficients to be equal to the highest and lowest values observed among open population, respectively. References for time since isolation (indicated by arrows) are reported in Supplementary Text S1.

### Population isolates in the European genomic background

Having described variation within populations, we next concentrated on their genetic relationships. We first explored the distance matrix based on inter-individual pairwise IBS distances. When sorting populations according to their average genetic distance, the highest value was found for Russians followed by Sauris, South Italy and Sicily (Fig. 6A). At the opposite end of the spectrum, the lowest values were observed for French, Basques, Lessinia Cimbrians and Aosta. Interestingly, the lowest pairwise genetic distances were recorded for the three mainland Sardinia populations, whereas among the German-speaking islands high levels of differentiation were found. Overall, we observed a significant correlation between the genetic and geographic distances considering the dataset both with (R2=0.209; p-value<0.05) and without (R2=0.203; p-value<0.05) the isolated populations (supplementary Figure S6). Even when applying a correction for the isolation-by-distance effect, patterns of open and isolated populations were found to be very similar (Fig. 6B).

**Figure 6.**
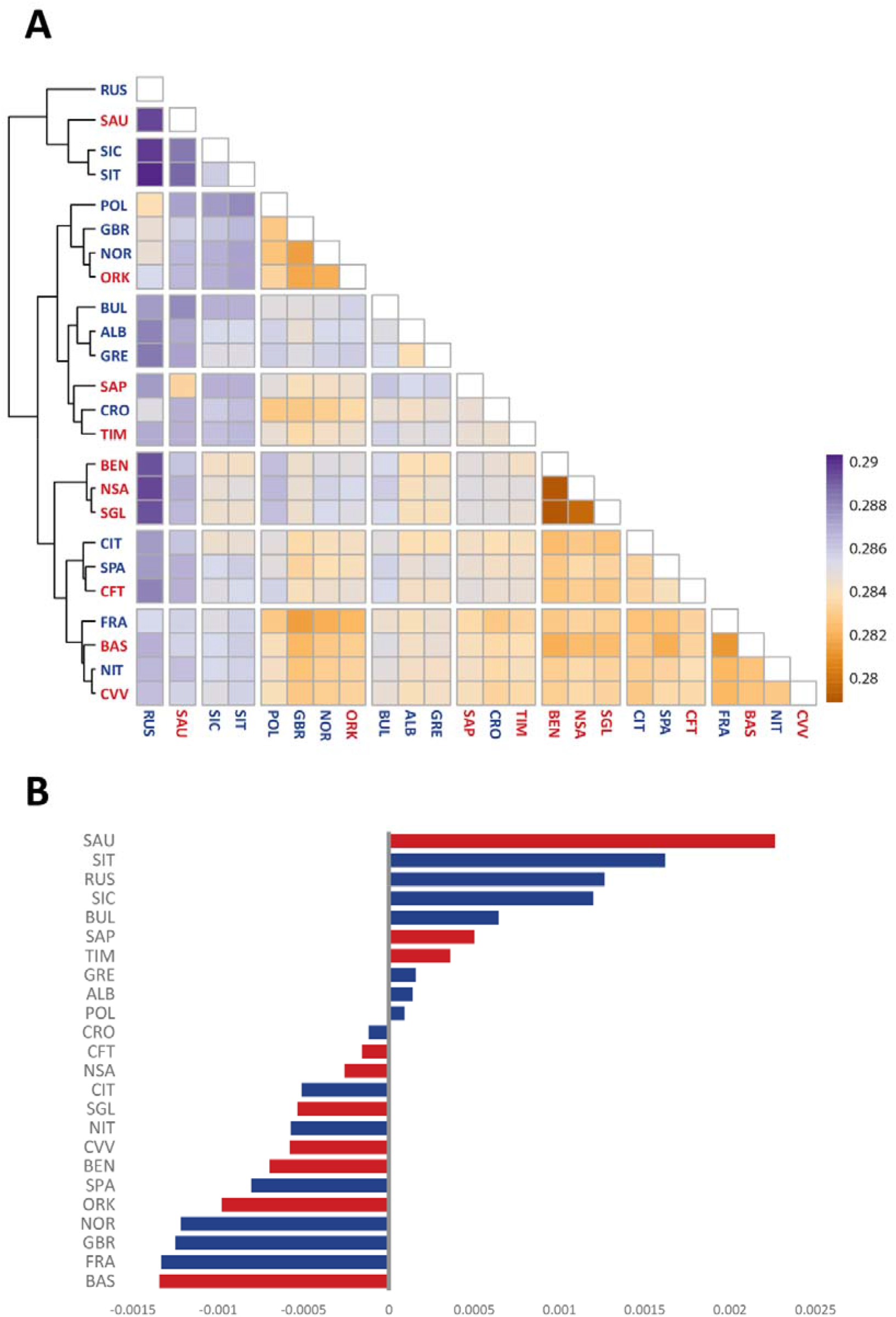
(A) Heatmap of pairwise genetic distances (R package Pheatmap). Populations are clustered according to a complete hierarchical approach (B) Deviation of the average genetic distances from those predicted by an isolation by distance model in open populations (see Materials and Methods for more details).

In order to capture more subtle signals of differentiation, we carried out a multidimensional scaling analysis (MDS)^43^. The first dimension clearly separates the three populations of mainland Sardinia (Benetutti, North Sardinia and Sulcis Iglesiente) from the others, whereas the second dimension distinguishes the two German-speaking islands of Sappada and Sauris (Fig. 7A). The third and fourth dimensions (Fig. 7B) separate the Basques (dimension 3) and Timau (dimension 4) respectively. These results recapitulate the PCA performed directly on SNP data (see Supplementary Figure S7). Furthermore, the best predictive model of ADMIXTURE (four ancestral populations; Supplementary Figure S8) reveals a pattern of population differentiation which is very close to that depicted by MDS.

**Figure 7.**
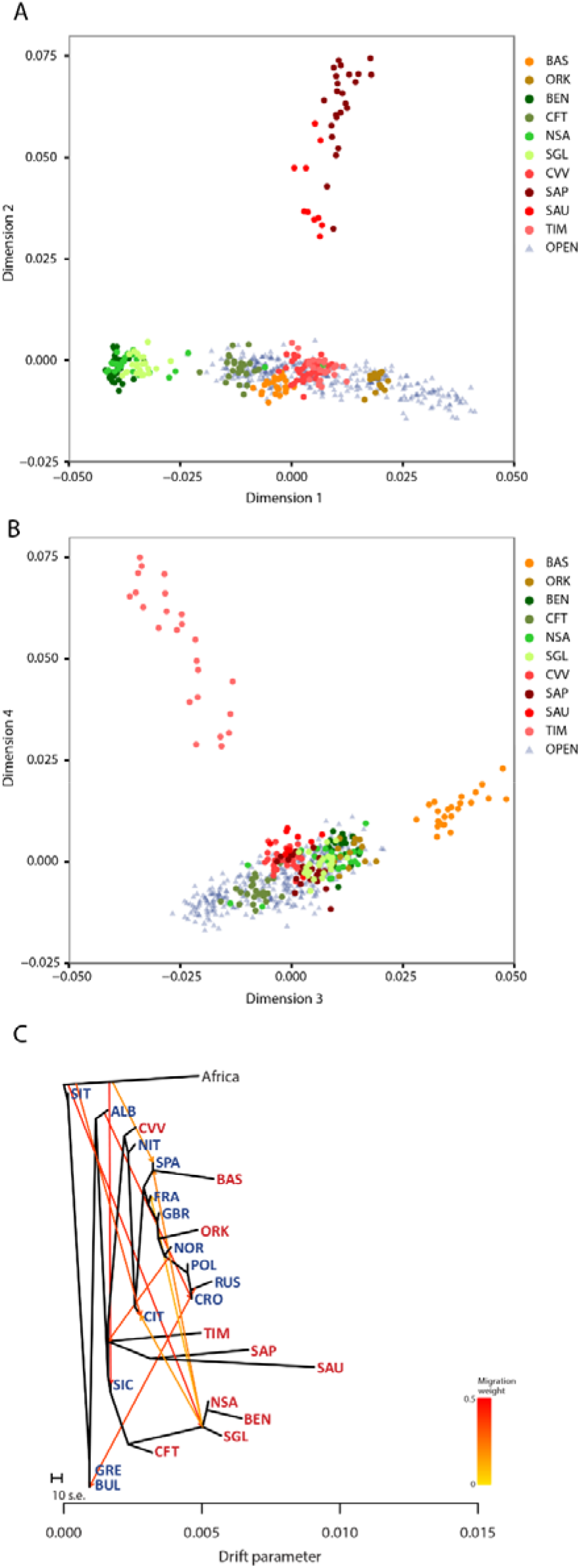
Plot of the first and second dimensions of the Multidimensional scaling analysis (A). Plot of the third and fourth dimensions (B). Treemix analysis with ten mixture events, with migration arrows coloured according to their weight (C).

Finally, we explored patterns of populations split and mixture using Treemix. The best fit with the data was obtained for the tree model with the maximum number of migration events tested, i.e. five (f=0.961). As shown in Fig. 7C, the general structure of the tree largely reflects the well-known relationships among European populations^34^. A high level of genetic drift can be observed for the entire Sardinian branch, and, even more pronounced for single German-speaking islands of Sappada, Sauris and Timau. Interestingly, the Cimbrians from Lessinia are more closely related to the Northern Italians and are located in the tree upstream to all northern and western European populations. Finally, although drifted away, Basques and Orcadians cluster on a geographic basis with Spain and close to British and Norwegians, respectively. Evidence of inward migration events involving our isolated populations was limited to the root of mainland Sardinians (from Africa), Sauris, Sappada and Timau (from a population ancestral to the Polish and Norwegians) and Basques (from the ancestral population of Sardinians).

## DISCUSSION

### New insights into the genomic diversity of isolated populations

All measures of intra-population genomic variation and the multivariate analysis (as visualized by the PCA plot) highlighted a relative homogeneity among European open populations, a finding which is in accordance with previous investigations^32,41,44^. However, the structure of genomic diversity was found to vary considerably among populations that have been subject to geographic and/or linguistic isolation. While such heterogeneity is consistent with what has been shown by gene-disease association studies^1^ and LD patterns^14^, our results may help shed more light on its extent and likely causes. We observed a greater dispersion of isolated than open populations along the first principal component, the variance of scores being 15.8 times higher for the former group. The scores were also found to be highly and significantly correlated with the inbreeding coefficient and drift parameter in the entire population dataset (total R2=0.901; p<0.01; see Supplementary Table S11 for further details). Although with a lower ratio (6.1), variation among isolates was higher also for the second principal component scores, which was found to be significantly correlated with effective size of isolates (R2=0.620, p-value=0.007). These results prompt a discussion of three different points.

Firstly, the principal component analysis helped disentangle the effects of the different forces that have shaped the genome of isolates. In fact, the analysis seems to indicate that most of their heterogeneity reflects variable intensities of drift and inbreeding, rather than their size or the time since isolation as suggested by historical sources. Taking the score of the first principal component as a tentative measure of isolation, Sauris, Sappada and Basques turned out to be the most isolated, while Orkney, Carloforte and Sulcis Iglesiente were the least, with Timau, Benetutti and North Sardinia in between.

Second, the fact we found populations with low scores for both the first and the second principal component - which means a combination of signatures of isolation with a relatively large effective size - calls into question the widespread view that human genetic isolates originate from a small group of founders^1,2,3,45^. The need for more complex models was earlier recognized by James V. Neel^46^, who proposed a categorization of isolates in which he included populations that originated from relatively large groups of individuals. Our study provides evidence that this idea is worth being further developed. In fact, whereas the estimates of current effective size for Sappada and Sauris seem not to contradict the idea of a small founding group, the values obtained for Basques and North Sardinia overlap with those estimated for a number of open populations. Whether or not this reflects a substantial difference in their founding population size should be considered with caution since our accuracy in estimating changes of effective population size over time^42^ was limited by the low SNP density of our genotyping platform. However, it should be noted that our study provides further indirect support to the view that population isolates may largely differ in the size of their founding groups^46^. In fact, when assuming demographic stationarity and equal effective population size among populations, the number of generations needed to reach the observed values of inbreeding coefficients was found to be substantially smaller for Basques and North Sardinians than for Sauris and Sappada (Supplementary Table S10 and Fig. 5). The evident discrepancy between these results and available knowledge about time since foundation of these isolates suggests that one or both assumptions are untenable. Although our findings are not sufficient to draw any definite conclusion, they show we must explore the extent of diversity in the size of founding groups of population isolates by using more powerful approaches based on whole genome sequences^47,48^.

Thirdly, our results suggest two quite distinct patterns of local isolation. In the case of the German-speaking islands, signals of heterogeneity among populations seem to prevail. Sappada, Sauris and Timau were found to be clearly different from each other both regarding intra and inter-population diversity. High genetic distances among Sauris, Sappada and Timau have already been observed with unilinear markers^9^, a pattern that is probably associated with the occurrence of a form of social behaviour which we termed “local ethnicity”. Despite their closely related languages and shared traditions, members of Alpine linguistic islands tend to identify their ancestry with their own village rather than considering themselves as part of the same ethnic group^9^. Such strong territoriality when defining ethnic identities and boundaries may have played a role in marriage strategies, decreasing the genetic exchange among the three linguistic islands. This “isolation among isolates” might have also allowed the genomic structure of each of them to evolve independently. On the other hand, a much greater homogeneity was observed among mainland Sardinians. The genetic distances among Benetutti, North Sardinia and Sulcis Iglesiente are the lowest in our dataset, and even lower than predicted by their geographic distances (Fig. 6B). This is not surprising because a similarity across the island has already been highlighted in previous studies^49,50,51^ (but see^52,53^). Therefore, despite the much longer time since isolation compared to German speaking islands, Sardinian populations seem to have maintained a certain homogeneity due to their larger effective population size which, in turn, could have weakened the effects of genetic drift and inbreeding. This could account for their lower variation of intra-population diversity measures, evidenced in the first principal component, relative to that observed among Sappada, Sauris and Timau.

### Continuity rather than dichotomy

Our analysis highlights a continuous pattern of genomic variation among populations that have been categorized as open and isolated. Looking at the first principal component, it is possible to identify a more dense area, which corresponds to the high homogeneity of open populations, and another more sparse zone, which reflects the greater diversity among isolates. In the contact zone, we found Cimbrians, who cluster along the first principal component with the open populations, while Carloforte and Sulcis Iglesiente are borderline. The behaviour of these three populations, all of which have been subject to both geographic and linguistic barriers in the course of their recent history, does not mean that their genomic structure is only marginally different from that of open populations. Rather, it points to the lack of clear discontinuities between the two groups when multiple indicators of isolation are used simultaneously. In fact, these three populations show signatures of past inbreeding which were undetected in the open ones.

This is effectively evidenced by their long upper tails of pairwise intra-population IBS and IBD block sharing (Supplementary Figure S4) and, in a more irregular fashion, by other measures of intra-population variation. A substantial continuity between open and isolated populations can be also appreciated looking at the PCA based on individual data, where only the 95% inertia ellipses of Sauris, Sappada and Basques do not overlap with that obtained by merging open populations (Supplementary Figure S9). A breakdown of the cultural barrier might account for the behavior of Cimbrians. In fact, only a limited number of individuals is today able to use the Cimbrian language^54^, a situation in contrast with the persistence of the original linguistic features in other German speaking communities^55^. This form of cultural assimilation, which started in the middle of the 16th century^56^, probably increased the permeability of Cimbrians to gene flow from neighbouring populations. Carloforte is the most recent isolate of our dataset, with the founding event dating back to 1738 AD. The small time since isolation and the genetic introgression associated with migratory waves from Tunisia, Liguria and Campania have presumably limited the effect of inbreeding and drift on the genome^57,58^. The attenuation of isolation signatures for Sulcis Iglesiente compared to other Sardinian populations may be explained by the more exogamous behaviour of villages of this area, a likely consequence of their location in coastal plains close to the Mediterranean Sea^59^.

At inter-population level, the picture obtained by using the genetic distance matrix directly does not discriminate between open and isolated populations. Previous studies revealed that diversity among European populations complies with an isolation by distance model on a continental^21,22^ and local scale^60,61^. We were able to find the same pattern over a wide continental range, regardless of the presence of isolates in the dataset, implying that they do not depart significantly from what is to be expected under isolation by distance. The only way to pinpoint a difference for some isolates was by considering specific MDS dimensions, which highlight a more pronounced scattering among individuals from Sauris, Sappada Timau and the Basques. Interestingly, these are also the populations in which we noticed the highest levels of inter-individual variation.

The overall picture provided by our study contrasts with previous observations based on unilinear markers. Using a dataset including all the isolates studied here, the only exception being the Orkney islands, we showed that most of them behave as outliers with both mtDNA (hypervariable region 1) and Y chromosome markers (six microsatellite loci)^62^. The discrepancy between genetic systems may be explained by the smaller effective size of unilinear markers, which makes them more prone to the effects of genetic drift, and the power of the above polymorphisms to detect also novel mutations that occurred after the population split.

## CONCLUSIONS

Through our study, we gain new insights into the genomic diversity of European populations that have been subject to linguistic and geographic barriers to gene flow. We were able to shed more light on their heterogeneity, challenge the generalized view of isolates as units that originated from small founding groups, and reveal that genomic patterns of open and isolated populations are distributed along a sort of continuum. We believe that there are two possible avenues to follow up these first results. Firstly, a comparison of the structure of open and isolated populations using whole genome sequences would provide a complete representation of their genomic diversity. Secondly, extending comparisons to geographical contexts other than Europe will help us understand to what extent the observed patterns may be appropriate to isolates in other continental or regional scenarios. Waiting for further investigations, we hope this first study can reach its own target: to make us more aware of the value of human population isolates in understanding how variable pressures of drift and inbreeding, determined by the interplay among environmental, socio-cultural and demographic factors, have shaped human genomic diversity.

## MATERIALS & METHODS

### Dataset

We assembled genome-wide SNP chip data of 561 healthy unrelated adult individuals from 24 European populations (Table 1 and Fig. 1). New genotype data were obtained for 211 subjects from three areas: (i) Sardinia (Benetutti, Carloforte, North Sardinia, Sulcis Iglesiente); (ii) German-speaking linguistic islands of the eastern Alps (Sappada, Sauris, Timau and Cimbrians from Lessinia); (iii) the Aosta province, in the Val d’Aosta region (north-western Italy). Only individuals with grandparents born in the same geographic area of sampling were enrolled in the study. Informed consent was obtained for all subjects. All methods were carried out in accordance with Italian Law (Decreto Legislativo della Repubblica Italiana, n° 196/2003). All experimental protocols were approved by the Bioethic Committee of the Azienda Ospedaliera Universitaria Pisana (Pisa, Italy. Prot N. 12702).

**Table 1.** Details on populations under study. N stands for sample size, G for Geographic and G/L for geo/linguistic. References for census size and time since isolation can be found in the Supplementary Text S1. Census sizes were obtained from the National population and housing census - 2011 (ALB, BEN, CIT, CFT, CRO, CVV, GBR, GRE, NIT, NSA, ORK, POL, SAP, SAU, SGL, SIC, SIT, SPA, TIM) – 2014 (BUL) – 2015 (RUS, NOR)-2016 (FRA).

| POPULATION | LABEL | N | CURRENT CENSUS | TIME SINCE ISOLATION<br>(in years before present) | ISOLATION FACTOR | REFERENCE |
| --- | --- | --- | --- | --- | --- | --- |
| **North Eastern Italian isolates** |  |  |  |  |  |  |
| Cimbrians | CVV | 33 | 13,455 | ~1000 | G/L | Present Study |
| Sappada | SAP | 24 | 1,307 | ~1000 | G/L | Present Study |
| Sauris | SAU | 10 | 429 | ~800 | G/L | Present Study |
| Timau | TIM | 24 | 500 | 800-1000 | G/L | Present Study |
| **Sardinian isolates** |  |  |  |  |  |  |
| Benetutti | BEN | 25 | 1,971 | ~5000 | G/L | Present Study |
| Carloforte | CFT | 25 | 6,301 | 268 | G/L | Present Study |
| North Sardinia | NSA | 25 | 96,448 | 3900-2900 | G/L | Present Study |
| Sulcis Iglesiente | SGL | 23 | 128,540 | 2800 | G/L | Present Study |
| **European isolates** |  |  |  |  |  |  |
| Orkney | ORK | 15 | 21,349 | ~1300 | G | 63 |
| French Basques | BAS | 24 | ~650,000<sup>64</sup> | 5500-3500 | G/L | 63 |
| **South Europe** |  |  |  |  |  |  |
| Albania (Gheg) | ALB | 24 | 2,831,741 | - | - | Sarno et al. in preparation |
| Croatia | CRO | 20 | 4,284,889 | - | - | 35 |
| Greece | GRE | 20 | 10,815,197 | - | - | 65 |
| Spain | SPA | 34 | 46,815,916 | - | - | 65 |
| **East Europe** |  |  |  |  |  |  |
| Bulgaria | BUL | 31 | 7,202,198 | - | - | 65 |
| Poland | POL | 32 | 38,511,824 | - | - | 65 |
| Russia | RUS | 25 | 144,192,450 | - | - | 63 |
| **North Europe** |  |  |  |  |  |  |
| Norway | NOR | 18 | 5,214,890 | - | - | 65 |
| British isles | GBR | 16 | 63,181,775 | - | - | 65 |
| **West Europe** |  |  |  |  |  |  |
| France | FRA | 28 | 67,264,000 | - | - | 63 |
| **Italy** |  |  |  |  |  |  |
| North Italy (Aosta) | NIT | 22 | 34,619 | - | - | Present Study |
| Central Italy (Piana di Lucca) | CIT | 25 | 394,318 | - | - | Tofanelli et al. unpub. data |
| South Italy | SIT | 18 | 14,184,916 | - | - | 65 |
| Sicily | SIC | 20 | 5,077,487 | - | - | 65 |

**Figure 1.**
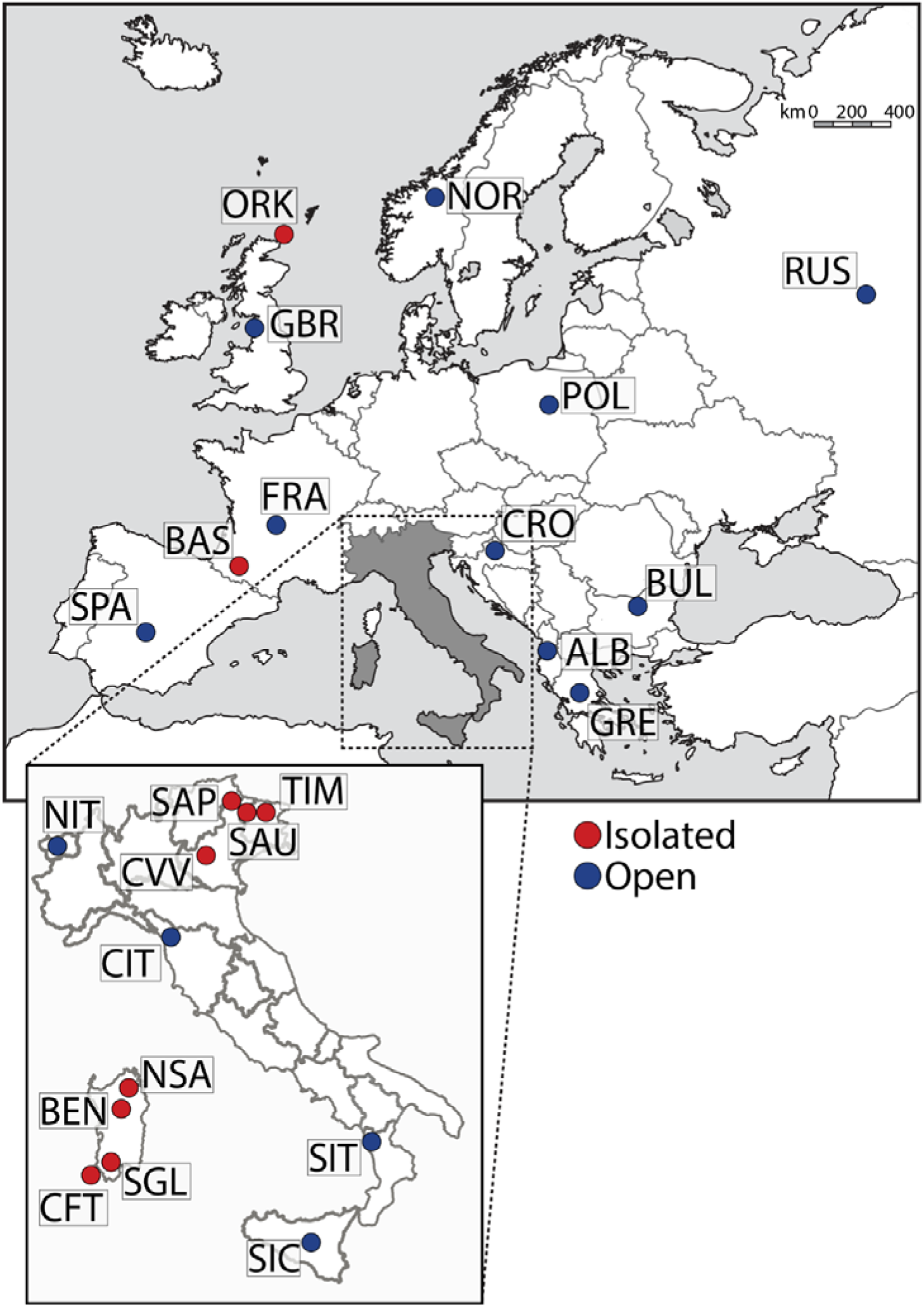
Map showing the geographic location of the 24 populations under study. Labels as in Table 1. Maps available from Wikipedia Common web page (https://commons.wikimedia.org/wiki/File:Blank political map Europe in 2006 WF.svg?uselang=it#fil elinks) were modified using Adobe Photoshop CS6 software.

The newly genotyped data were analyzed jointly with literature data for a total of 10 isolated and 14 open populations (Table 1). Concerning the choice of isolates, our dataset included populations with historically documented barriers to gene flow (geographic and linguistic) and a broad range of census size and time since isolation (see Table 1; see Supplementary Text S1 for a detailed description of populations under study). This was coherent with our aim to explore the diversity occurring among isolates - defined inclusively as populations which have been subject to geographical and/or cultural factors of isolation during their evolutionary history - avoiding any bias due to arbitrary selection. Regarding the selection of open populations, we considered the following three criteria: (i) geographic proximity with the isolated population dataset; (ii) geographic coverage of the European continent; (iii) sample size of at least 15 individuals.

### Genotyping, quality control and validation

All samples were genotyped using the Geno 2.0 DNA Ancestry Kit (www.genographic.com) SNP microarray known as the GenoChip^24^ at the Gene-by-Gene laboratory (Family Tree DNA) in Houston, Texas. The autosomal AIMs (Ancestry Informative Markers) implemented in the GenoChip array provide an adequate coverage of the genetic diversity of European populations^24^, and include rare variants occurring in small sized population samples. Furthermore, the geographic homogeneity of the typed populations minimized confounders that could potentially have originated from ascertainment bias when performing cross-population comparisons. The newly genotyped samples were merged with the reference ones and then filtered according to the standard genotype quality control metrics using PLINK^25^. Only the SNPs with a genotyping success rate > 90% were included, giving a total of 87,818 autosomal SNPs after the addition of the literature data. Only the individuals with a genotyping success rate > 92% were used. Relatedness to the 3rd generation (Identity by Descent, IBD > 0.185) was tested with PLINK, and from the detected relative pairs, only one sample was randomly chosen for the subsequent analysis. We tested the power of the set of selected SNPs in detecting signals of isolation comparing them with those contained in the HGDP panel (647,789 SNPs) (see Supplementary Text S2).

### Intra-population analyses

Runs of homozygosity (RoHs) were estimated using PLINK v1.9 (--homozyg option) (https://www.cog-genomics.org/plink2) under default settings (sliding window of 5 Mb, minimum of 50 SNPs, one heterozygous genotype and five missing calls allowed). Each SNP was considered to be part of a homozygous segment when the proportion of overlapping homozygous windows is above 5%. RoHs were defined as stretches of at least 0.5 Mb with at least 25 homozygous SNPs^17^. We performed an unsupervised Gaussian fitting of the length distribution using *Mclust* from the R package mclust V3 ^26^ and identified six different classes based on RoH’s length (Supplementary Figure S1).

Pairwise shared identical-by-descent (IBD) segments were identified by the refined IBD algorithm implemented in Beagle v.4.1^27^ adopting default parameters. Thereafter, we used the pairwise length of IBD sharing to calculate the statistic W_int_^28^. This index represents the total length of the shared IBD blocks averaged over the number of possible pairs of individuals.

IBS values were estimated using PLINK v1.9 (--distance ibs option). By default, this option produces a lower-triangular tab-delimited text file with pairwise IBS between all individuals in the dataset. From this matrix, we extracted the values calculated between pairs of individuals belonging to the same population in order to obtain the intra-population pairwise IBS.

Blocks in linkage disequilibrium were calculated by using --blocks option (default settings) in PLINK v1.9. We used the -hardy option in PLINK v1.9 to obtain the average observed heterozygosity (het) per population and the inter-locus variance, calculated as the square root of the standard deviation. Homozygosity was calculated as hom=(1-het).

The Levene test for the equality of variances was performed with the R software package Car^29^.

Principal component analysis^30^ was performed with the R software package Ade4^31^ using the above-described intra-population measures as variables.

Multiple regression analysis was performed with R software using the scores of the first and second principal component as dependent variables and the inbreeding coefficient, drift parameter (inferred by Treemix, considering the value from the nearest tree node, see below), the effective population size point estimates and the time since isolation as independent variables. The inbreeding coefficient was calculated as the proportion of the autosomal genome in runs of homozygosity, excluding the centromeres^32^.

### Inter-population analyses

Maximum likelihood estimation of individual ancestries was performed using ADMIXTURE vl.23^33^ under default values (the block relaxation algorithm, a termination criterion set to stop when the log-likelihood increases by less than ε = 10^−4^ between iterations and the quasi-Newton convergence acceleration method with q = 3 secant conditions). We applied unsupervised clustering analysis to the whole sample set, exploring the hypothesis of K=1 to 10 clusters. We assessed cross-validation errors for each value of K using the ADMIXTURE’S Cross Validation procedure.

MDS analysis was performed using PLINK v1.9 (–distance-matrix option). The information carried by each dimension was assessed by calculating the ratio of their respective eigenvalues compared to the sum of all eigenvalues.

Genetic structure and gene flow were investigated using TreeMix v1.1^34^. We set the position of the root (root option) using a North African population (Egyptians^35^). To account for the fact that nearby SNPs are not independent, we grouped them together in windows of 500 SNPs using the -k flag. We used the -m option to build the maximum likelihood tree with one to five migration events to minimize data overfitting, and we then calculated the fraction of the variance in relatedness between populations explained by each model.

Geographic distance matrix was calculated using the Geographic Distance Matrix Generator (http://biodiversitvinformatics.amnh.org/opensource/gdmg). The geographical coordinates of the sampling areas were downloaded from the http://maps.cga.harvard.edu/gpf/. When the exact locations of sampling were unknown, we used the coordinates of the centroid of the nation as reported in http://gothos.info/resources/. In order to control for the effects of geographical proximity, we calculated the deviation of any observed genetic distance from the one predicted by the regression line obtained for geographic and genetic distances of open populations.

### Estimate of the effective population size

Effective population size for all populations was estimated using the LD method^36^ implemented in NeEstimator 2.0^37^. The LDNe algorithm estimates effective population size from the extent of linkage disequilibrium in the sample. Pairwise LD was calculated between 68,205 autosomal SNPs from 16 randomly chosen chromosomes. Other random combinations gave results that were very close to those reported in Supplementary Table S2. We decided against using the entire dataset because with too many loci, the method could not compute confidence intervals. We used a threshold of 0.05 as the lowest allele frequency, which gives the least biased results^38^. We reported estimated N_e_ with 95 % (parametric) confidence intervals.

The number of generations since isolation and the relative values of effective population size under a model of demographic stationarity were calculated from the formula^39^,

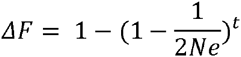

where *ΔF* stands for the difference between the inbreeding coefficient estimated for each population and its hypothetical value at the time of population split. The latter parameter was assumed to range between the highest and lowest inbreeding coefficients observed among open populations.

## ACKNOWLEDGEMENTS

We are greatly indebted to all the blood donors. We would also like to thank Marcella Benedetti (Municipality of Sappada), Nino Pacilè and Lucia Protto (Municipality of Sauris), Vito Massalongo (Giazza), Ottaviano Matiz and Velia Plozner (Timau) for their valuable assistance in sample collection and for their warm hospitality.

This study was supported by a 2013 National Geographic Society Genographic 2.0 grant to ST. The survey in the Eastern Italian Alps was also funded by the Università di Roma “La Sapienza” (ref. C26A13HSHB) and the Istituto Italiano di Antropologia. SS, AB and DP were supported by the European Research Council ERC-2011-AdG 295733 grant (Langelin).

## AUTHOR CONTRIBUTIONS STATEMENT

Conceived and designed the study: GDB PA ST VD. Provided samples: GDB CC ST DP GV. Extracted and prepared DNA for genomic analysis: CB CC ST. Analyzed the data: PA VD LP ST PF VC SS AB. Contributed reagents/materials/analysis tools: ST GDB LP MGV RSW. Wrote the manuscript: GDB PA in collaboration with LP and VC. Read and approved the final manuscript: all authors.

## ADDITIONAL INFORMATION

**Accession codes** (where applicable)

All data are available at the Zenodo Database (https://zenodo.org/) with accession number: 10.5281/zenodo.50114.

## Competing financial interests

The authors declare no competing financial interests.

